# Targeting CXCR4-expressing TAMs in muscle-invasive bladder cancer to enhance tumor control after immunotherapy

**DOI:** 10.1101/2025.03.21.644543

**Authors:** Emma Desponds, Hajar El Ahanidi, Nagham Alouche, Hana Zdimerova, Stéphanie Favre, Giulio Zanette, Sina Nassiri, Daniel Benamran, Petros Tsantoulis, Mohammed Attaleb, Hélène Maby-El Hajjami, Julien Dagher, Vanessa Gourhand, Karl Balabanian, Marion Espeli, Sanjiv A. Luther, Camilla Jandus, Marine M Leblond, Grégory Verdeil

## Abstract

Bladder cancer (BC) is a prevalent malignancy with poor prognosis in advanced stages. While immune checkpoint blockade has revolutionized immunotherapy, its efficacy remains limited for most advanced BC patients. The detailed characterization of BC’s tumor microenvironment (TME) is a prerequisite to understand these mechanisms of resistance and to develop new therapeutic strategies. In this study, we used a genetically engineered BC mouse model resistant to anti-PD1 treatment, and BC patient samples, to investigate the evolution of tumor-associated macrophages (TAMs) during BC progression. We identified a subset of pro-tumor TAMs expressing CXCR4, predominantly found in advanced stages of BC-bearing mice and in half of muscle-invasive BC patients from the studied cohort. Interestingly, CXCR4^+^ TAM-rich regions were associated with CD8 T cell-excluded areas in both mice and patients. Administration of a small molecule CXCR4 inhibitor significantly reduced the number of pro-tumor TAMs within the tumor and markedly prolonged mouse survival. Incorporating this inhibitor into a tri-immunotherapy regimen further enhanced survival, highlighting the potential of targeting multiple pathways to strongly enhance anti-tumor effects and offering new hope for improving immunotherapy in advanced BC.

## Introduction

Bladder cancer (BC) represents a prevalent health concern, ranking as the ninth most common cancer worldwide with approximately 220’000 deaths annually (1). At diagnosis, about 75% of BC patients are identified with non-muscle invasive BC (NMIBC), while the remaining 25% faces muscle invasive BC (MIBC) (2). Treatments for this latter stage offer limited effectiveness as the 5-year survival rate is around 50% (3), posing a major challenge in the field of uro-oncology. Despite the high tumor mutational burden of BC (4), only 20% of MIBC patients respond to programmed cell death protein 1 (PD-1)/programmed death ligand 1 (PD-L1) inhibitors (5). Immune checkpoint inhibitors (ICIs) combined with the antibody-drug conjugate (ADC), Enfortumab-vedotin, have shown survival improvement. However, the median of progression-free survival is only 12 months (6), emphasizing the importance of finding additional targets to further improve patients’ outcome.

The tumor microenvironment (TME) is known to play a critical role in tumor progression and treatment resistance (7), but the TME of BC remains poorly investigated compared to other solid tumors. Tumor-associated macrophages (TAMs) are one of the key players of the TME. TAMs can display various functions ranging from supporting tumor growth to exerting anti-tumor effects (8) and different subsets can co-exist in the same tumor (9). In BC, TAMs are the most abundant tumor-infiltrating immune cell population (10) and are frequently associated with unfavorable clinical outcomes and treatment response (11). While appealing, completely deleting TAMs can be counterproductive in some cases, as some immunotherapies depend on anti-tumor TAMs for effective tumor control (9, 12–14). However, targeting the right TAM populations at the appropriate moment is still an unsolved challenge. For the moment, TAMs-targeting therapies, such as CSF1R inhibition, have demonstrated limited efficacy in clinics in various cancer types, accentuating the need to identify new appropriate TAM targets.

The chemokine CXCL12 and its receptors CXCR4/CXCR7 were already described to play an important role in the development of various tumors (15). In BC, mRNA expression for both *CXCL12* (16–18) and *CXCR4* (19) were associated with poor survival. While several studies focused on the expression of CXCR4 and CXCR7 on bladder tumor cells (20–25), few have investigated the role of CXCR4 in the TME. Interestingly, omics analyses of BC tissues have shown a positive correlation between *CXCL12* gene expression and the presence of immunosuppressive TAMs (17, 26, 27). Moreover, several clinical trials in various cancer types have reported that CXCR4 inhibition indicated encouraging clinical efficacy, especially in combination with other treatments (28). Despite these findings, the therapeutic potential of targeting TAMs via the CXCL12-CXCR4 pathway remains unexplored in BC.

In this study, we investigated the evolution of MHCII^low^ and MHCII^high^ TAMs, previously classified as pro- and anti-tumor TAMs, respectively (9, 29, 30), throughout BC progression in a genetic mouse model of MIBC replicating key aspects of the human pathology (9). We identified a subset of pro-tumor TAMs that expressed CXCR4, which was mostly found in the advanced stages of BC-bearing mice. In patients, half of the MIBC patients’ samples studied were infiltrated with CXCR4-expressing TAMs and correlated with low CD8 T cell infiltration. Pharmacologic inhibition of CXCR4, by itself, was sufficient to improve the survival of MIBC-bearing mice. Furthermore, we demonstrate a synergistic effect of ICIs and adjuvant CXCR4 inhibition in MIBC, that can be beneficial for CXCR4^+^ TAM-infiltrated patients to improve ICI treatment’s efficacy.

## Results

### TAMs evolve toward a pro-tumor phenotype along with bladder tumor progression

We previously reported the conversion from an anti-tumor towards a pro-tumor TME along the NMIBC to MIBC transition in an inducible mouse model of BC (9). To understand in more detail the mechanism behind this evolution, we analyzed TAMs during tumor progression. We observed that most TAMs are MHCII^high^ in NMIBC while they shift towards an MHCII^low^ pro-tumor phenotype when tumor invades the muscle layers (Fig.1A). To define more accurately TAMs evolution through BC progression, RNA sequencing analyses were performed on isolated MHCII^low^ and MHCII^high^ macrophages in healthy bladder and at the different stages of the disease (Suppl. Fig.S1.A). A two-dimensional projection revealed that MHCII^low^ and MHCII^high^ are distinct populations of macrophages (Fig.1.B). Moreover, for both phenotypes, TAMs from NMIBC are closely related to the macrophages from healthy bladder while TAMs from both muscle-invasive stages clustered together (Fig.1.B). When comparing MHCII^low^ and MHCII^high^ macrophages across stages, MHCII^high^ macrophages expressed higher level of pro-inflammatory associated genes, such as *Ccl5*, *Cxcl9*, *Cxcl10*, *Cxcl11*, *Il-1β*, *Il-12*, *Ifnβ*, and *Ifnγ*, while MHCII^low^ macrophages expressed pro-tumor genes, such as *Msr1*, *Retnlb*, *Mmp9*, *Tgfβ2*, *Il-4* and *Xpr1* (Fig.1.C) (31). As tumors progressed, gene expression signatures from both MHCII^high^ and MHCII^low^ macrophages evolved, each population showing dynamic changes in specific sets of genes (Fig.1.D). During tumor progression, MHCII^high^ macrophages exhibited a decrease in pro-inflammatory genes expression (e.g., *Ccl5*, *Il-6*, *Ccl8*, *Cd40*, *Ifnβ* and *Ifnγ*), while gaining immunosuppressive traits (e.g., *Tgfβ3*, *Mmp9* and *Arg1*) (Fig.1.D; Suppl. Fig.S1.B). In parallel, MHCII^low^ macrophages increased the expression of angiogenic genes (e.g., *Angpt2*, *Vegfc* and *Vegfβ*), pro-tumor chemokines (e.g., *Cxcl3* and *Cxcl5*) and anti-inflammatory cytokines (e.g., *Mif* and *Tgfβ3*) with tumor progression (Fig.1.D; Suppl. Fig.1.B). To find whether TAMs’ population were also evolving in patients, we analyzed TAMs in fresh NMIBC and MIBC patients’ samples. Flow cytometry analyses showed that TAMs (defined as CD14^+^HLA-DR^+^ cells - Suppl. Fig.S1.C), and especially pro-tumor CD163^+^ TAMs [29], were more abundant in MIBC compared to NMIBC (Fig.1.E,F). To confirm this result and localize TAMs in the bladder tissue in a larger cohort of MIBC patients, we stained paraffin-embedded sections for CD68 and CD204 from 26 MIBC patients. We showed that pro-tumor CD204^+^CD68^+^ macrophages [30] were mainly contained in tumor cores (Fig.1.G,H) and increased especially in pT3 and pT4 stages (Fig.1.I). Moreover, TCGA analysis showed that a high pro-tumor TAM signature (31) was associated with poor overall survival in MIBC (Fig.1.J). Overall, the data in BC-bearing mice demonstrate a gradual evolution of macrophage populations during disease progression towards acquiring pro-tumor gene signatures. Similarly, higher numbers of TAMs expressing pro-tumor markers were also found in human advanced BC stages, correlating with worse survival.

**Figure 1.**
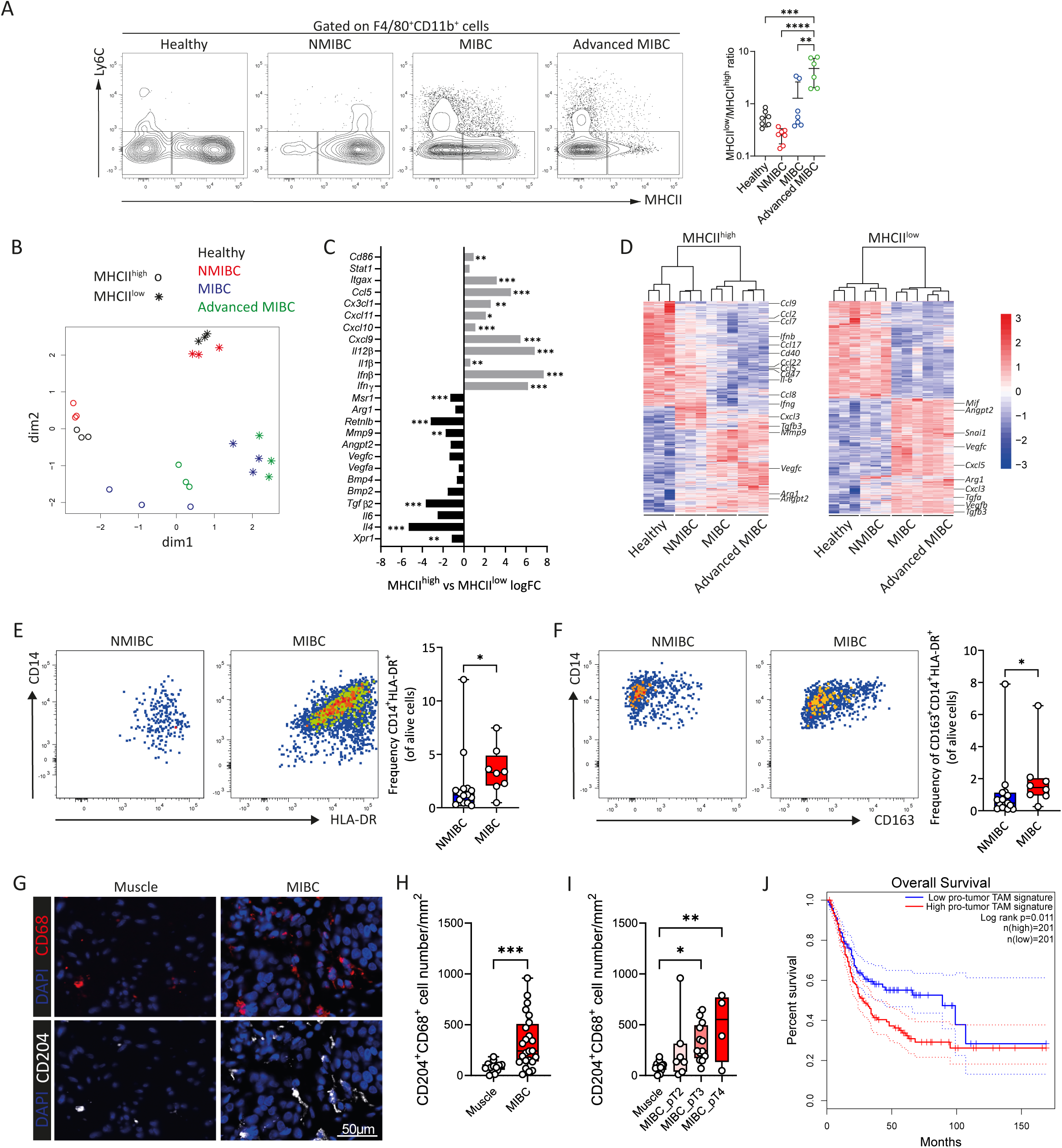
Evolution of macrophage populations with bladder tumor progression. (**A**) Representative flow cytometry contour plot of Ly6C and MHCII expression on macrophages with the corresponding histogram of the MHCII^low^ and MHCII^high^ macrophage ratio in healthy, NMIBC-, MIBC- and advanced MIBC-bearing mice. Each dot represents an individual mouse, and bars represent the mean ± SD. One-way ANOVA, followed by Tukey’s HSD test. (**B**) Multidimensional scaling of gene expression of sorted MHCII^high^ and MHCII^low^ macrophages in healthy, NMIBC-, MIBC- and advanced MIBC-bearing mice. Each dot represents an individual mouse. (**C**) Differentially expressed genes between sorted MHCII^high^ versus (vs) MHCII^low^ macrophages from our mouse model of BC with all-time points pooled. (**D**) Heatmap of differentially expressed genes in sorted MHCII^high^ and MHCII^low^ macrophages in healthy, NMIBC-, MIBC- and advanced MIBC-bearing mice. The data is row-wise standardized using z-scores. (**E-F**) Representative flow cytometry dot plots and frequency of CD14 and HLA-DR expressions (**E**) and CD14 and CD163 expressions (**F**) in NMIBC and MIBC patients. Each dot represents an individual patient. Nonparametric Mann-Whitney test. (**G**) Representative CD68 (red), CD204 (white) and DAPI (blue) immunofluorescent images of human bladder tumor cores and adjacent muscle of MIBC patients. (**H**) Quantification of CD204^+^CD68^+^ cell number in muscle and MIBC. Each dot represents an individual patient. Nonparametric Mann-Whitney test. (**I**) Quantification of CD204^+^CD68^+^ cell number in adjacent muscle and in pT2, pT3 and pT4 MIBC. Each dot represents an individual patient. One-way ANOVA, followed by Tukey’s HSD test. (**J**) Kaplan-Meier survival analysis of pro-tumoral TAM signature in MIBC patients based on TCGA data.

### The CXCR4-CXCL12 axis promotes pro-tumor macrophage accumulation during bladder cancer progression

To identify potential targets limiting the accumulation of pro-tumor TAMs during BC progression, we screened for chemokine and cytokine ligand-receptor pairs. Transcriptomic analysis of the whole bladder in healthy-, NMIBC-, MIBC- and advanced MIBC-bearing mice revealed several potential pathways upregulated in advanced MIBC, including *Cxcr4-Cxcl12*, *Csf1-Csf1r* and *Ccr2-Ccl2* (Fig.2.A). To validate these results, we measured levels of 15 chemokines in the bladder supernatant and urine obtained from mice at the different stages of the disease (Fig.2.B; Suppl. Table S1). CXCL12 was the only chemokine with increased level at MIBC stages in both bladder supernatant and urine (Fig.2.B). When staining bladder slides at the various stages of the disease, we found that CXCL12, both mRNA and protein, were faintly detectable in healthy tissues but significantly increased in MIBC stages (Fig.2.C). To decipher a potential involvement of these pathways on TAM biology in BC, we studied the protein expression of the related receptors at the different stages of BC. Both CCR2 and CSF-1R were mainly expressed on MHCII^high^ TAMs and their expression decreased with tumor progression (Suppl. Fig.S2.A). Conversely, CXCR4 was specifically expressed on MHCII^low^ TAMs and showed an increased expression level at advanced stages (Fig.2.D), when CXCL12 protein’s expression significantly increased (Fig.2.C). These CXCR4^+^ TAMs had higher expression of Arg1, CD204, CD206 and PD-L1 than CXCR4^-^ TAMs (Fig.2.E), which represents a stronger pro-tumor phenotype signature. In MIBC-bearing mice, TAMs were the major populations expressing CXCR4 in tumors (Suppl. Fig.S2.B), while Ly6C^high^ monocytes have the highest CXCR4 expression in blood (Suppl. Fig.S2.C). To test whether CXCR4 expression on macrophages is regulated by the TME, tumor conditioned media (TCM) from advanced stages was incubated with bone marrow derived macrophages (BMDMs). TCM promoted CXCR4 expression on these BMDMs (Suppl. Fig.S2.D). Altogether, our findings suggest a potential role of the CXCR4-CXCL12 pathway in the accumulation of pro-tumor TAMs in advanced BC.

**Figure 2.**
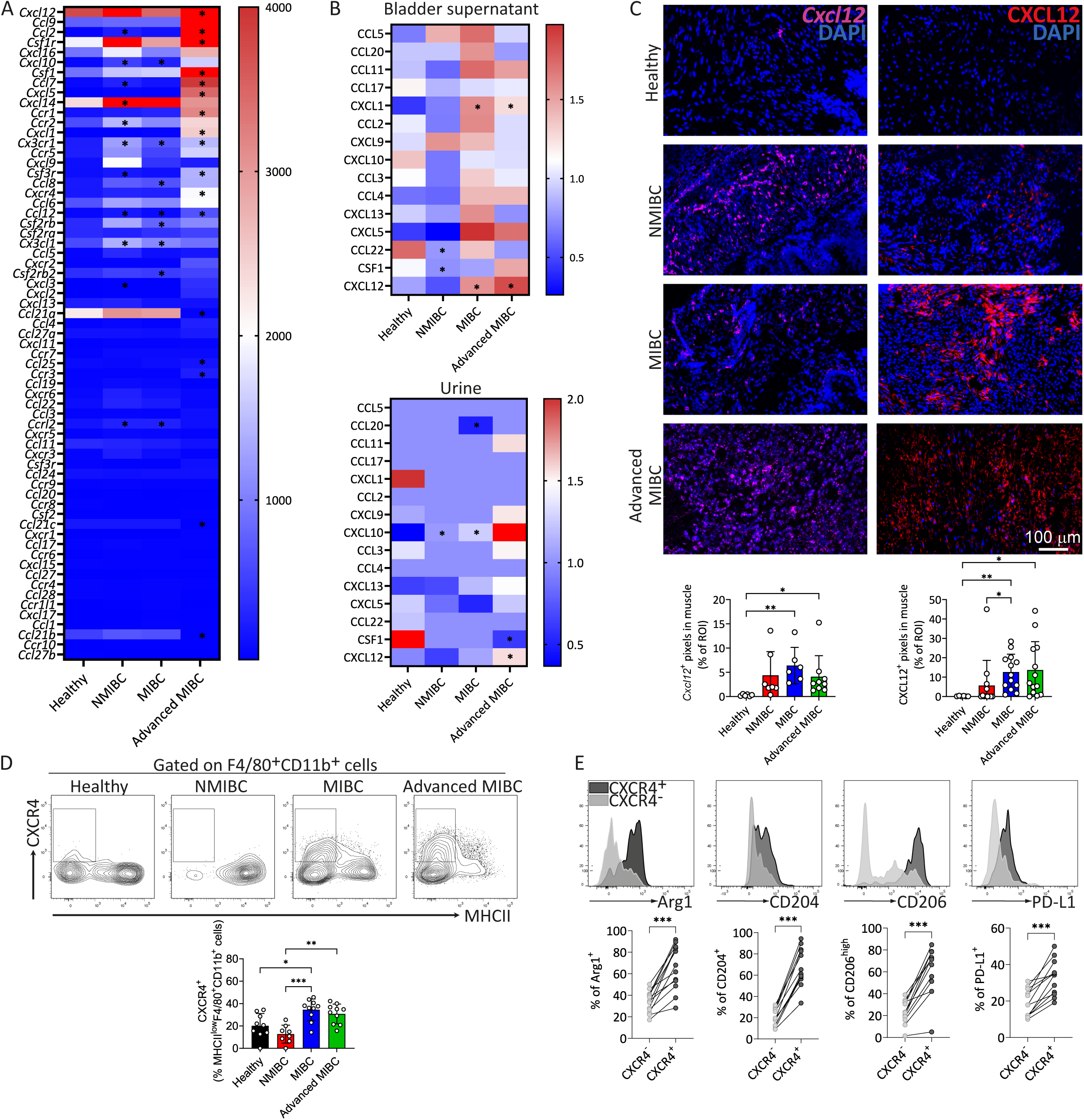
Evolution of CXCL12 expression and CXCR4^+^ macrophage infiltration during bladder tumor progression in mice. (**A**) Heatmap of the mean RNA count for the selected chemokines and cytokines in the whole bladder of healthy, NMIBC-, MIBC- and advanced MIBC-bearing mice. (**B**) Heatmap of chemokine concentration normalized to the median of each chemokine in bladder supernatant and urine from healthy, NMIBC-, MIBC- and advanced MIBC-bearing mice. (**C**) Representative immunofluorescent images of DAPI (blue), *Cxcl12* RNA (pink) and CXCL12 protein (red) of bladder from healthy, NMIBC-, MIBC- and advanced MIBC-bearing mice. Quantification of the percentage of CXCL12^+^ pixels of the ROIs for mRNA and protein analyses. Each dot represents an individual ROI, and bars represent the mean ± SD. One-way ANOVA, followed by Kruskal-Wallis test. (**D**) Representative flow cytometry contour plots of CXCR4 and MHCII expressions on macrophages in healthy, NMIBC-, MIBC- and advanced MIBC-bearing mice and the corresponding bar plot with percentage of CXCR4^+^ macrophages. Each dot represents an individual mouse, and bars represent the mean ± SD. One-way ANOVA, followed by Tukey’s HSD test. (**E**) Representative flow cytometry histograms and dot plots of the frequencies of Arg1, CD204, CD206 and PD-L1 expression on CXCR4^+^ and CXCR4^-^ TAMs in MIBC- and advanced MIBC-bearing mice. Each dot represents an individual mouse. Wilcoxon matched-pairs signed rank test.

### CXCL12 expression and infiltration of CXCR4^+^ pro-tumor TAMs increased in advanced stages of human BC

To confirm the clinical relevance of the CXCR4-CXCL12 pathway for patients, we analyzed CXCR4 and CXCL12 levels in human BC samples. Using the TCGA dataset, we observed that the mRNA expression of both *CXCR4* and *CXCL12* increased with the BC staging (Fig.3.A). While high levels of *CXCL12* correlated with poorer overall survival in human MIBC, *CXCR4* expression did not (Fig.3.B). At the protein level, CXCL12 concentrations were higher in the serum, but not in the urine, of BC patients compared to healthy individuals (Fig.3.C). Flow cytometric analysis of fresh human BC tissues revealed an increased frequency of CXCR4^+^ TAMs in MIBC compared to NMIBC, with a high heterogeneity in the MIBC group (Fig.3.D). As observed in preclinical settings, the phenotype of these CXCR4^+^ TAMs showed typically increased expression of pro-tumor markers such as CD163, Arg1, Tie2 and CD204 (Fig.3.E). Immunofluorescence staining on a cohort of 26 MIBC patients’ samples showed that the number of CXCR4^+^CD68^+^ macrophages increased in the tumor core compared to non-tumor muscle for 53,8% of patients (14 out of 26 patients, Fig.3.F,G), especially pT3 and pT4 tumors (3/7 pT2 patients; 9/13 pT3 patients; 2/4 pT4 patients; Fig.3.H). Moreover, in this cohort of patients, CXCR4 staining in the tumor core is mostly co-localized with the CD68 staining (Fig.3.F), suggesting that CXCR4 is preferentially expressed by TAMs. These results confirmed in patients that CXCR4^+^ pro-tumor TAMs increased in advanced stages of bladder tumors and could affect patients’ survival.

**Figure 3.**
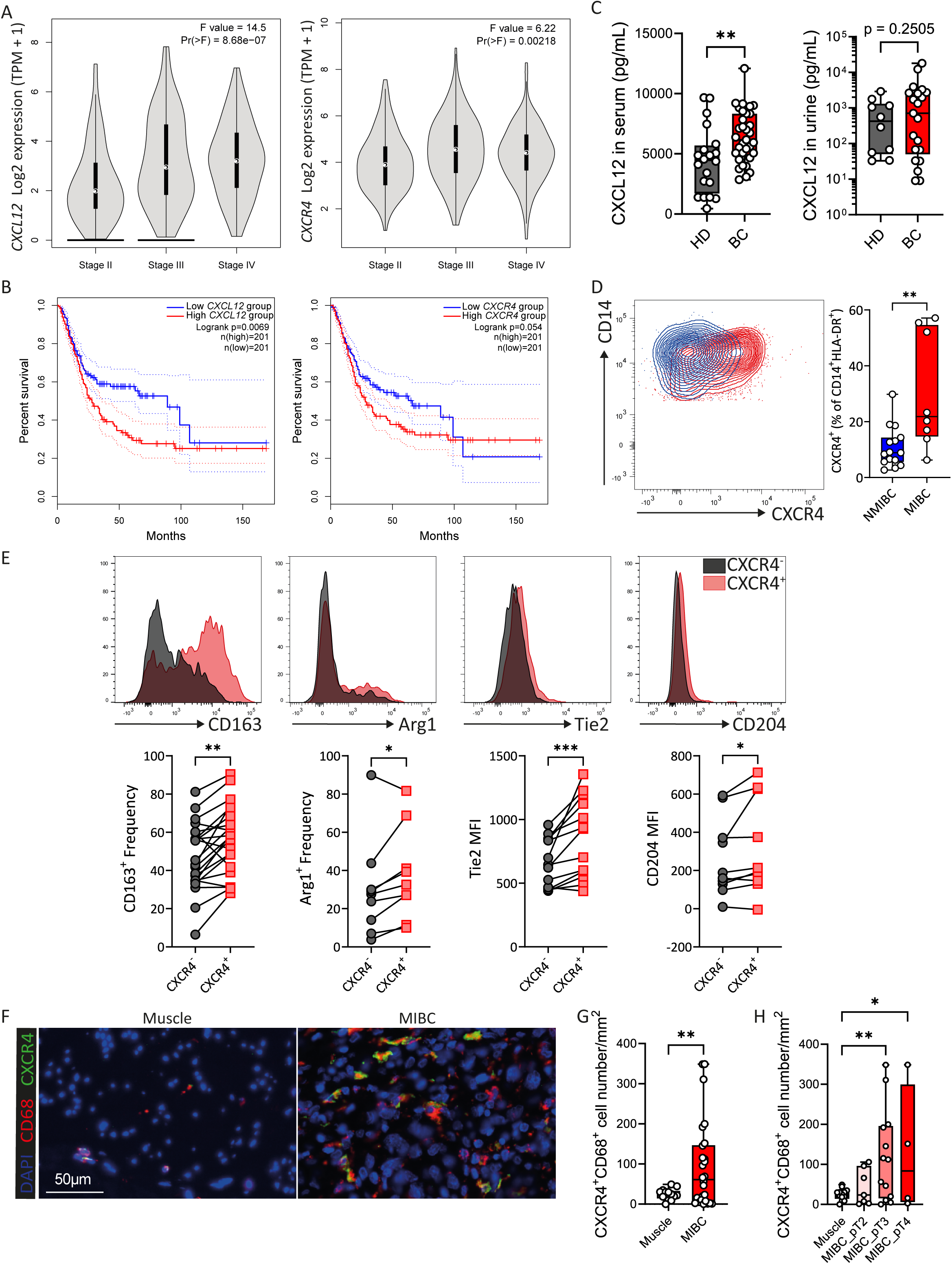
CXCL12 and CXCR4 expressions in human bladder cancer. (**A**) Violon plot of *CXCL12* and *CXCR4* expression over stage II, stage III and stage IV from MIBC patients, as based on the TCGA datasets. (**B**) Kaplan-Meier overall survival analysis of *CXCL12* and *CXCR4* expression in MIBC patients based on TCGA data. (**C**) Box plots of CXCL12 concentration in the serum and urine of healthy donor (HD) and BC patients. Nonparametric Mann-Whitney test. (**D**) Representative flow cytometry contour plot of CD14 and CXCR4 expressions on macrophages in NMIBC and MIBC patients and the corresponding bar plot with percentage of CXCR4^+^ macrophages. Each dot represents an individual patient. Nonparametric Mann-Whitney test. (**E**) Representative flow cytometry histograms of indicated markers on CXCR4^+^ and CXCR4^-^ macrophages from BC patients and their corresponding frequency or mean fluorescent intensity (MFI) dot plots. Each dot represents an individual patient. Wilcoxon matched-pairs signed rank test. (**F**) Representative CD68 (red), CXCR4 (green) and DAPI (blue) immunofluorescence microscopy images of MIBC patients. Scale bar=50 µm. (**G**) Quantification of CXCR4^+^CD68^+^ cell numbers in muscle and MIBC. Each dot represents an individual patient. Nonparametric Mann-Whitney test. (**H**) Quantification of CXCR4^+^CD68^+^ cell number in muscle and in pT2, pT3 and pT4 MIBC. Each dot represents an individual patient. One-way ANOVA, followed by Tukey’s HSD test.

### CXCR4^+^ TAMs-infiltrated muscle-invasive bladder cancers display a CD8 T cell-excluded environment

Since we previously noticed that the increase of pro-tumor TAMs paralleled the decrease in CD8 T cells in mice (9), we investigated the relationship between CXCL12 levels, CXCR4^+^ TAMs and CD8^+^ T cell infiltration in BC. In mice, we observed that a high infiltration of CXCR4^+^ TAMs was found in CXCL12^+^ areas, which coincided with a low infiltration of CD8 T cells (CD8b^+^CD3^+^ cells) (Fig.4.A). Conversely, MIBC with low/no CXCL12 had low/no infiltration of CXCR4^+^ TAMs but high infiltration of CD8 T cells (Fig.4.A). In patients, we mentioned previously that MIBC patients could be divided in two categories, i.e., the ones with high infiltration of CXCR4^+^CD68^+^ cells (>100 cells/mm^2^; 7/13 pT3 patients; 2/4 pT4 patients) or the ones with low frequency of CXCR4^+^CD68^+^ cells (<100 cells/mm^2^; 6/13 pT3 patients; 2/4 pT4 patients) (Fig.3.F,G,H). We confirmed in patients that CXCR4^+^ TAMs were also mainly localized in CXCL12-rich areas (Fig.4.B,C), while CD8^+^CD3^+^ cells were localized predominantly in CXCL12^-^ areas (Fig.4.B,D). Interestingly, CXCR4^+^ TAMs infiltration was negatively correlated with the infiltration of CD8 T cells (Fig.4.E). Overall, these results indicate that CXCR4 is expressed on TAMs in approximately half of advanced MIBC patients and that MIBC patients with high infiltration of CXCR4^+^ TAMs displayed very low numbers of CD8 T cells. On the other hand, MIBC patients without CXCR4^+^ TAMs infiltration have a higher infiltration of CD8 T cells in their tumor.

**Figure 4.**
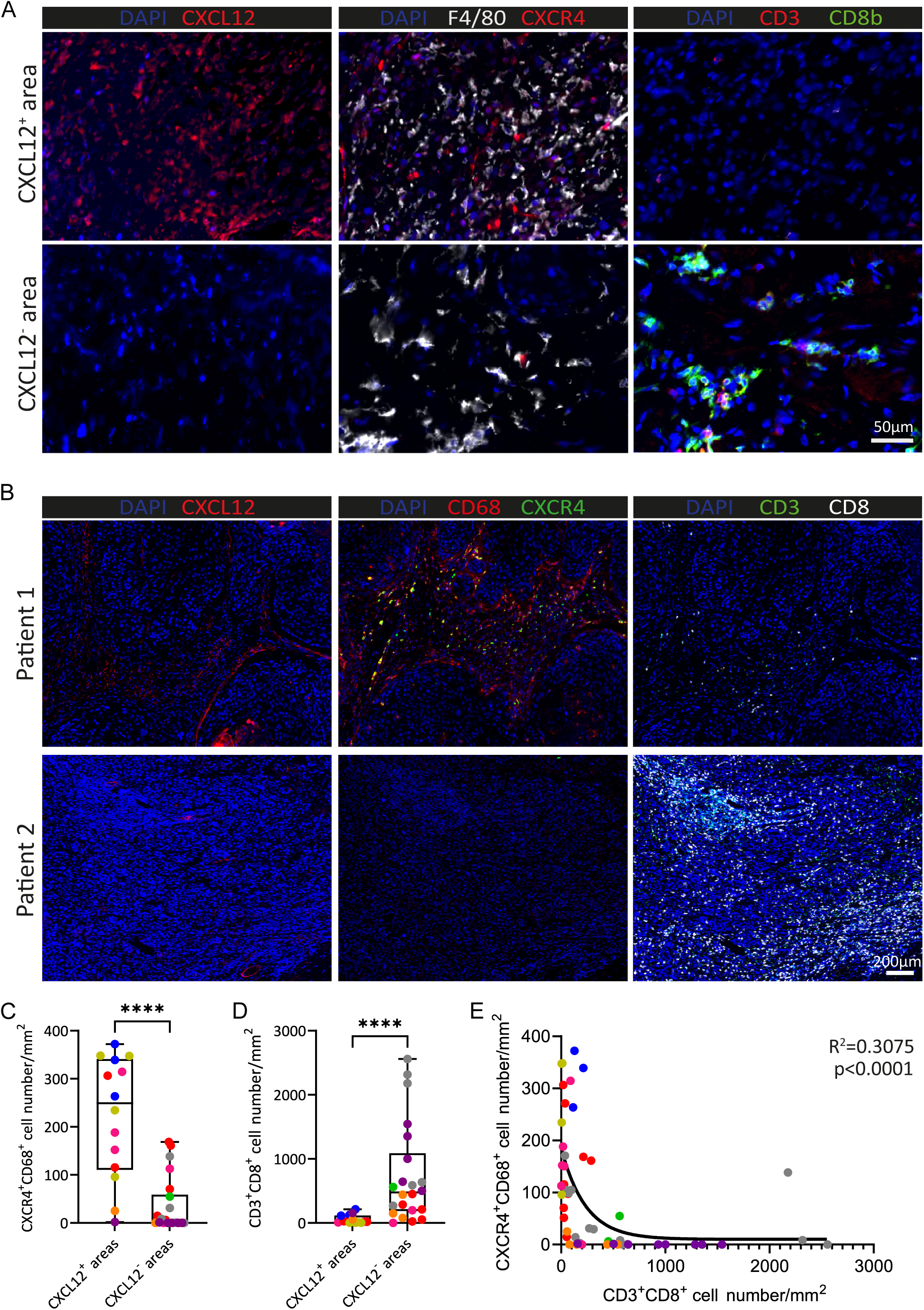
Localization of CXCR4^+^ TAMs and CD8 T cells in bladder cancer. (**A**) Representative CXCL12, F4/80, CXCR4, CD3, CD8b and DAPI immunofluorescent images of MIBC-bearing mice. Scale bar=50 µm. (**B**) Representative CXCL12, CD68, CXCR4, CD3, CD8 and DAPI immunofluorescent images of MIBC patients with high infiltration of CXCR4^+^ TAMs (patient 1) or low infiltration of CXCR4^+^ TAMs (patient 2). Scale bar=200 µm. (**C-D**) Quantification of CXCR4^+^CD68^+^ cell numbers (**C**) and CD3^+^CD8^+^ cell numbers (**D**) in CXCL12^+^ and CXCL12^-^ areas from MIBC patients. Nonparametric Mann-Whitney test. (**E**) Correlation between the number of CXCR4^+^CD68^+^ cells and CD3^+^CD8^+^ cells in MIBC patients. Two phase decay nonlinear regression analysis with a nonparametric Spearman correlation. Quantifications were performed on at least three regions of interest for four MIBC patients with high infiltration of CXCR4^+^ TAMs and four MIBC patients with low infiltration of CXCR4^+^ TAMs. Each dot represents an individual region of interest, and each color represents a single patient.

### CXCR4 blockade decreases pro-tumor TAMs and increases survival of mice with MIBC

To assess the therapeutic potential of CXCR4 blockade in an anti-PD-1 resistant model of MIBC, we used the small molecule inhibitor, AMD3100, that specifically antagonizes CXCR4 (32). AMD3100 was administrated in drinking water of MIBC-bearing mice throughout this mouse survival experiment (Fig.5.A). AMD3100 treated mice showed a significant increase in survival of approximately 26 days compared to the control group (Fig.5.B), while CSF-1R or CCL2/CCR2 TAM-targeting treatments did not (Suppl. Fig.S3.A,B). Nine days post-treatment with AMD3100, we don’t observe a decrease in bladder weight (Fig.5.C), but we noticed a decrease in the number of the MHCII^low^ TAM subset without any effect on the MHCII^high^ subset compared to the control group (Fig.5.D). AMD3100 treatment did not impact the number of Ly6C^high^ monocytes, CD8 T cells, conventional CD4 T cells or regulatory T cells in the bladder (Fig.5.E) nine days after the beginning of treatment. However, a reduction in Ly6G^+^ cell number was noticed nine days post-treatment (Fig.5.E), although these cells did not express CXCR4 (Suppl. Fig.S2B). As AMD3100 treatment did not increase MHCII^high^ TAMs in the tumor, nor did it impact macrophage proliferation or survival in vitro (Fig.5.F), we suspect that AMD3100 blocks the recruitment of MHCII^low^ TAM. Despite the significant delay of survival after ADM3100 treatment alone, the mice ultimately succumbed to BC. To further improve survival, we considered combining CXCR4 blockade with other immunotherapies. Therefore, we tested the combination of CD40 agonist and PD-1 blocking antibodies (9) with AMD3100 in MIBC-bearing mice (Fig.5.G). The tri-therapy significantly improved mouse survival compared to control group or the CD40 agonists and anti-PD-1 bi-therapy, but only when AMD3100 was administrated in an adjuvant regimen, but not in a concomitant regimen (Fig.5.G). Altogether, these results demonstrate that CXCR4 blockade efficiently decreases pro-tumor TAMs in MIBC and acts in synergy with immunotherapies targeting antigen-presenting cells and T cell activation, such as CD40 agonists and anti-PD-1 treatments, respectively.

**Figure 5.**
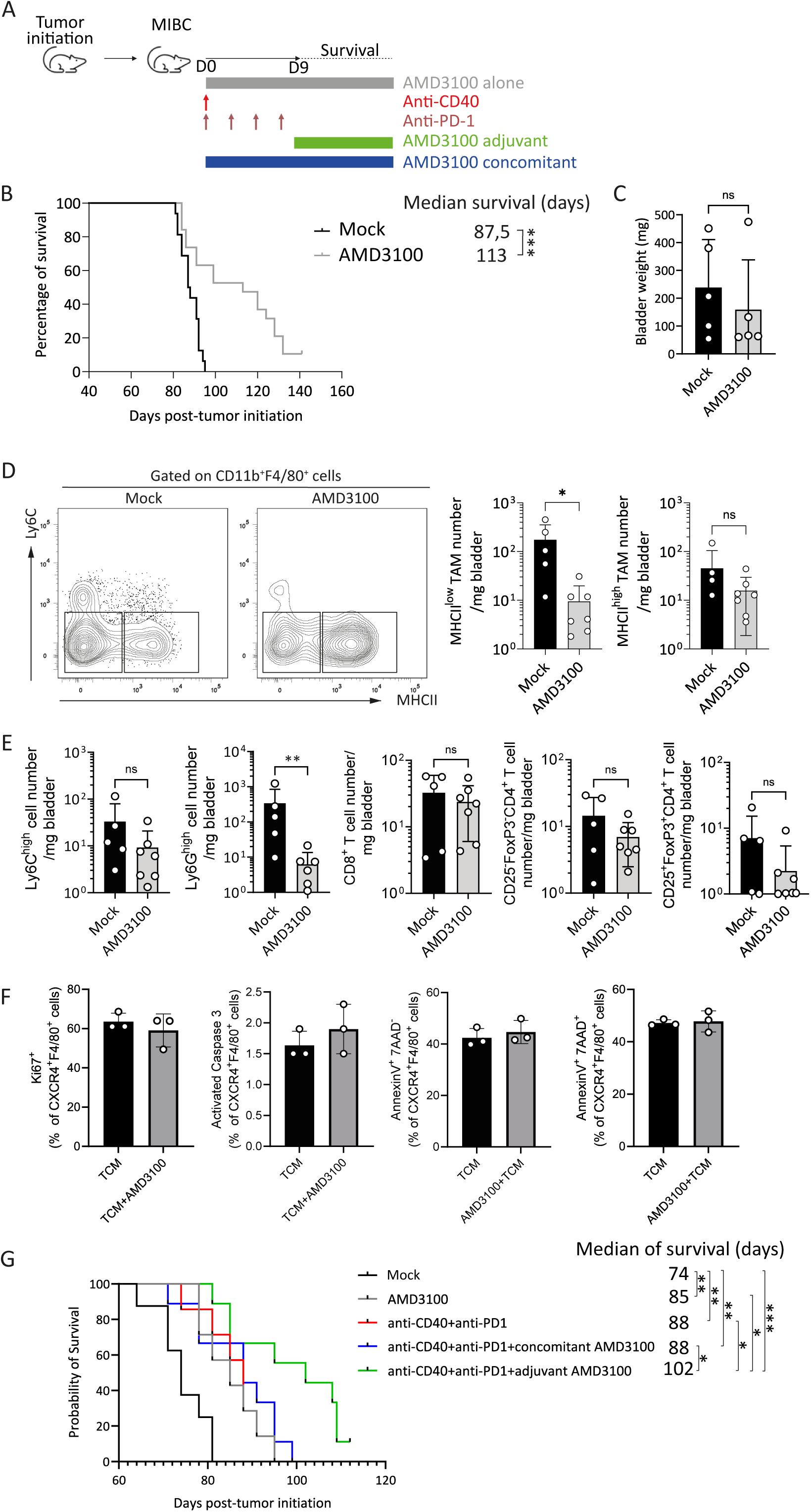
Effects of CXCR4 blockade in MIBC-bearing mice. (**A**) Timeline of AMD3100 treatments, anti-CD40 agonist and anti-PD-1 blocking antibody in MIBC-bearing mice. AMD3100 alone was administrated at D0, concomitant AMD3100 was administrated at the same time as anti-CD40 and anti-PD-1 on D0, adjuvant AMD3100 was administrated at the end of anti-PD-1 treatment on D9. (**B**) Kaplan-Meier curve of MIBC-bearing mice treated with AMD3100 or water as control group (Mock). n=15. Log-rank test. (**C**) Bladder weight of MIBC-bearing mice treated with AMD3100 or water as control group (Mock). Each dot represents an individual mouse, and bars represent the mean ± SD. Nonparametric Mann-Whitney test. (**D**) Representative flow cytometry contour plot of Ly6C and MHCII expressions on macrophages in bladders of control (mock) or AMD3100 treated MIBC-bearing mice with the quantification of MHCII^low^ and MHCII^high^ macrophage numbers. Each dot represents an individual mouse, and bars represent the mean ± SD. Nonparametric Mann-Whitney test. (**E**) Quantification of Ly6C^high^, Ly6G^high^, CD8^+^ T cell, CD25^-^FoxP3^-^CD4^+^ T cell and CD25^+^FoxP3^+^CD4^+^ T cell numbers in the bladder of mock MIBC-bearing mice or treated with AMD3100. Each dot represents an individual mouse, and bars represent the mean ± SD. Nonparametric Mann-Whitney test. (**F**) Bar plots showing frequencies of Ki67, activated caspase 3, annexinV^+^ 744D^-^ and annexinV^+^744D^+^ bone marrow-derived macrophages treated with tumor conditioned medium (TCM) or TCM+AMD3100. Each dot represents an individual experiment, and bars represent the mean ± SD. (**G**) Kaplan-Meier curve of MIBC-bearing mice treated with isotype control antibody (Mock), AMD3100, anti-CD40+anti-PD-1, anti-CD40+anti-PD-1+concomitant AMD3100 or anti-CD40+anti-PD-1+adjuvant AMD3100. n=8. Log-rank test.

## Discussion

The development of macrophage-targeting therapies has opened new perspectives for cancer treatment. However, the diverse states of TAMs and their rapid ability to adapt their phenotype within the TME complicate the development of therapies that specifically target the pro-tumor subtypes. Previous studies on the inhibition of the colony-stimulating factor 1 receptor (CSF-1R) (33, 34) or of the CCL2/CCR2 axis (35, 36) led to promising results in preclinical studies of other types of cancer. In BC-bearing mice, CSF1-R and CCR2 were not associated with pro-tumor TAMs and blocking these pathways did not improve mouse survival, suggesting that targeting these pathways is inefficient for the treatment of MIBC. We identified a pro-tumor TAM population expressing CXCR4, which was found in advanced stages of BC-bearing mice, but also in half of the studied MIBC patients. While most BC studies investigate the role of CXCR4 on tumor cells, especially in metastasis development (20–24, 37), our research offers a different perspective by focusing on CXCR4 expression in macrophages. In mice and in the studied cohort of MIBC, we detected most of CXCR4 expressions extracellularly on TAMs. Various reasons could explain these discrepancies, such as the antibody used, the conditions of the staining or the lack of proper identification of macrophages versus tumor cells in immunohistology. We do not rule out a potential role of CXCR4 on tumor cells, but we established a clear correlation between CXCR4^+^ TAM presence, CD8 T cell infiltration and BC progression.

The pharmacological inhibition of CXCR4 led to a significant decrease of MHCII^low^ TAMs in MIBC-bearing mice, in accordance with other preclinical studies on hepatocellular carcinoma (38) or breast cancer (39). Interestingly, CXCR4 blocking did not affect the MHCII^high^ anti-tumor TAMs. An even more interesting strategy would consist in reeducating TAMs toward an anti-tumor phenotype (33) to provide a more tumor-restrictive environment. Despite that, the decrease of MHCII^low^ TAMs in the tumor after CXCR4 blockade paralleled a decrease in neutrophil. Knowing that this cell type does not express CXCR4 in MIBC-bearing mice, this suggests that the effects of CXCR4 blockade may primarily impact only pro-tumor TAMs, which in turn could regulate neutrophil accumulation, increasing in this way the immunosuppressive TME at the latest stage of BC in mice (9).

Inhibiting CXCR4 at the muscle-invasive stage was accompanied by a significant increase of survival in mice, without noticeable side effects. While our study is the first to block CXCR4 in a mouse model of MIBC, comparable results were observed in a study focusing on the NMIBC stage (37). As NMIBC patients from our cohort have lower infiltration of CXCR4-expressing TAMs compared to those with more advanced disease, targeting CXCR4 might be more beneficial in the advanced stages of the disease. Additionally, the high heterogeneity in CXCR4^+^ TAM infiltration within MIBCs suggests that patients might respond differently to CXCR4 blockade, with a better response in patients displaying high infiltration. In our cohort of MIBC, CXCR4^+^ TAM-infiltrated tumors were associated with low CD8 T cell infiltration and vice-versa. This suggests that patients with high infiltration of CXCR4^+^ TAMs will respond less to ICIs than patients with a low infiltration of CXCR4^+^ TAMs. Thus, selecting patients based on CXCR4 expression levels may represent an interesting new avenue to adjust the therapeutic management of MIBC.

To test potential synergistic effects of CXCR4 blockade with other therapies, we tested a multiple-therapy approach. Limiting TAMs accumulation with CXCR4 blockade following radiotherapy has already been investigated at the clinical level (40). Alternatively, trials also investigated the combination of CXCR4 blockade with other immunotherapies (41). In clinical settings, the combination of CXCR4 blockade with anti-PD-1 treatment has shown modest yet promising anti-tumor efficacy in solid tumors, such as advanced melanoma (42) or renal cell carcinoma (43). Interestingly, we observed a significant improvement of mouse survival with our combo-therapy compared to monotherapies or the combination of CD40 agonist and anti-PD-1 alone only when the CXCR4 inhibitor was administrated as an adjuvant with CD40 agonist with anti-PD-1. As already described, CD40 agonist and anti-PD-1 requires MHCII^high^ TAMs to induce an efficient anti-tumor effect (9). We suspect that blocking too early CXCR4 after CD40 agonist and anti-PD-1 treatment reduces TAM reeducation toward an anti-tumor phenotype, while blocking CXCR4 at the end of the bi-therapy allowed the reversal of the TME and then prolong survival by inhibiting the recruitment of new pro-tumor TAMs. This highlights the importance of deciphering the mechanism of action of each specific treatment to determine the most efficient sequence of administration, as already described in different tumors for immunochemotherapy (44–46).

To adapt patient treatment management as fast as possible, biomarkers that can predict patient response are urgently needed. In BC, PD-L1’s association with ICI response remains controversial (11, 47). Alternative biomarkers that are more specific and less invasive are therefore of great medical importance. Based on the findings of our current study, we speculate that CXCR4 expression could be used as a biomarker to orientate therapeutic management of MIBC patients. While CXCR4 expression was mainly detected by histology on biopsies (21, 22), innovative imaging techniques have been explored to detect CXCR4 by non-invasive imaging techniques, involving coupling a CXCR4 antagonist molecule with either a radionuclide or a fluorochrome (21, 48). However, these techniques require specific equipment and the injection of radio-labelled molecules into the patients. Non-invasive blood-based signatures that can predict patient response are an emerging area in the field of biomarker discovery (49–51). We showed in this study that CXCR4 expression was the highest on monocytes in the blood of MIBC-bearing mice. Moreover, we previously reported that *Cxr4* mRNA expression decreased in the blood of mice treated after CD40 agonist and anti-PD-1 combo-therapy (52), indicating its potential for predicting therapeutic response. However, a deeper analysis to determine whether mice with the highest CXCR4 expression in blood respond the least to ICIs would further strengthen its predictive potential.

Altogether, we demonstrate in this study the scientific rationale for targeting pro-tumor TAMs through CXCR4 inhibition, both in mice and in MIBC patients. We showed that blocking CXCR4 decreased pro-tumor TAMs in MIBC and achieved significant improvement in mouse survival in combination with other immunotherapies. This approach provides new insights for improving immunotherapeutic strategies, patient selection and MIBC outcomes.

## Methods

### Bladder cancer patient samples

Fresh blood and tumor samples were obtained from BC patients from the University Hospital of Geneva in the frame of the study protocol n° 2020/02375 approved by the Commission cantonale d’éthique de la recherche sur l’être humain (CCER), canton of Geneva, and upon written informed consent. Patients’ clinical information is summarized in Supplementary Table S2. Serum was extracted from blood prior surgery. Tumor samples were excised by surgeons in the frame of trans-urothelial resection of the bladder tumor (TURBT) or during cystectomy. Tumor biopsies were freshly collected in Leibovitz’s L-15 medium (# 11415064, ThermoFisher Scientific), supplemented with Hepes (#15630080, ThermoFisher Scientific), Glucose (#49163, Merck) and penicillin-streptomycin (# 15140122, ThermoFisher Scientific). Small tumor pieces were cut and digested in complete Leibovitz’s L-15 medium containing 2.5mg/ml of Liberase^TM^ (#5401119001, Merck) for 30 minutes at 37°C. For big tumor pieces, tumors were dissociated in complete Leibovitz’s L-15 medium containing 2.5mg/ml of Liberase^TM^ using the gentleMACS™ dissociator instrument (Miltenyi Biotec) and gentleMACS™ C tube (#130-093-237, Miltenyi Biotec), following the manufacturer’s protocol. Cell suspension was filtered using a 70µm strainer and red blood cells lysis was performed using red blood cell lysis Buffer (Qiagen). Cells were then ready to be stained for flow cytometry analysis or were cryopreserved.

Urine samples from BC patients were obtained in the frame of the study protocol N82/19, approved by the Ethics Committee for Biomedical Research from the Faculty of Medicine and Pharmacy of Rabat-Morocco, upon written informed consent. Urine samples were centrifuged (3500 rpm, 15 minutes) and supernatant was collected and cryopreserved. Patients’ clinical information is summarized in Supplementary Table S3.

### Mouse model of BC

All animal experiments were performed in compliance with the University of Lausanne Institutional regulations and were approved by the veterinarian authorities of the Canton de Vaud (authorizations VD3430, VD3594 and VD3856). Tp53^Fl/Fl^Pten^Fl/Fl^ mice were obtained by crossing Tp53^Fl/Fl^ mice (B6.129P2-Trp53tm1Brn/J) with Pten^Fl/Fl^ mice (B6.129S4-Ptentm1Hwu/J) purchased from Jackson Laboratories. To induce bladder tumors, 2.5×10^8^ plaque-forming units of Cre-expressing adenoviral vector [#AVL(VB181004-1095pzc)-K1, VectorBuilder, USA] in 5µl of DMEM/hexadimethrine bromide (8mg/ml) was injected into the bladder lumen of Tp53^Fl/Fl^Pten^Fl/Fl^ mice by micro-surgery as already described (53).

### Therapeutic treatments

Treatment schedule is available in Figures 5A and S3A. Briefly, therapeutic treatments started eight weeks after vector injection, when tumors reached the muscle-invasive stage and were palpable, and mice were sacrificed at nine days after the beginning of treatments to analyze the immune microenvironment or left for monitoring survival. Anti-PD1 blocking Ab (300µg/dose, RMP1-14 clone, BioXcell), or IsoCT (300µg/dose, 2A3 clone, BioXcell), was injected by i.p. injections every two-three days for eight days, as already published (9). Mice received one i.p. injection of anti-CD40 Ab (100µg/dose, FGK45 clone, BioXcell), or IsoCT (100µg/dose, 2A3 clone, BioXcell). Anti-CSF1R blocking Ab (600µg/dose, AFS98 clone, BioXcell), or IsoCT (600µg/dose, 2A3 clone, BioXcell), was injected once per week by i.p. injections. Anti-CCL2 blocking Ab (100µg/dose, 2H5 clone, BioXcell), or IsoCT (100µg/dose, InVivoMAb polyclonal Armenian hamster IgG, BioXcell), was injected daily by i.p. injections. CCR2 inhibitor (2mg/dose, PF-4136309, MedChemExpress) was injected subcutaneously daily. CXCR4 inhibitor (AMD3100, 3299/50, Bio-Techne) was used at 60µg/ml in drinking water. AMD3100 solution was changed every two-three days.

### Single-cell preparation from mouse blood and tissue

Erythrocytes from blood were eliminated with red blood lysis buffer before staining. Healthy and tumor bladders were first digested for 30min at 37°C in complete RPMI (RPMIc, 10% FCS, 1% penicillin/streptomycin), 0.1mg/ml DNase I (#D4527, Sigma), 1mg/ml Collagenase I (#17100017, ThermoFisher Scientific). Tissues were then mashed through a 70µm cell strainer. To isolate leukocytes from bladders, samples were centrifuged in density gradients 40%/70% Percoll for 30min at 2000 rpm. Isolated cells were washed in RPMIc before staining.

### Supernatant preparation from mouse bladders and tumor cells

Bladder supernatant from healthy or tumor bladders was obtained by putting the whole bladder in complete DMEM (DMEM, 10% FCS, 1% penicillin/streptomycin) at 40mg of tissue per ml. Supernatant was collected after 24h of incubation at 37°C.

To isolate tumor cells, single cell suspensions were performed as described above. Tumor cells were isolated by centrifugation in density gradients 75%/100% Percoll (#17-0891-01, GE Healthcare Life Sciences) for 30min at 2000rpm. Isolated cells were washed and put in culture in RPMIc. Tumor cell supernatant was obtained by collecting supernatant after 24 hours.

### In vitro assays with BMDMs

Isolation of BMDMs were performed by flushing the femurs and tibias of mice with IMDM (IMDM, 60% FCS and 1% penicillin–streptomycin). The cells were then washed and platted in IMDM (IMDM, 15% FCS, 1% penicillin–streptomycin) with 10ng/ml of Flt3-Ligand and 10ng/ml of M-CSF. The cells were cultured for 7 days before being used in an in vitro assay. BMDMs were plated into 24-well plate in 500µl of tumor condition media (TCM) from advanced bladder tumors in the presence or absence of AMD3100 (1µg/ml, #3299/50, Bio-Techne).

### Flow cytometry staining and analysis

FcγR were blocked for 15min at RT with αCD16/32 (1/1000, #101320, Biolegend). After staining for extracellular markers and then viability using LIVE/DEAD™ Fixable Aqua Dead Cell Stain Kit (#L34966, ThermoFisher Scientific) or the Zombie NIR Fixable Viability Kit antibody (#423106, Biolegend), cells were fixed and permeabilized with the Foxp3 Transcription Factor Staining Buffer Set (#00-5523-00, eBiosciences) according to manufacturer’s instructions. Intracellular staining was performed in permeabilizing buffer. To detect active Caspase 3, cells were cultured for 4 hours at 37°C. After extracellular staining, cells were stained intracellularly with a primary anti-Caspase 3 antibody for 1 hour at 4°C. Following a wash, a secondary antibody was added and incubated for 15 minutes. For Annexin V staining, the cells were initially stained on the surface and then processed using the Annexin V-APC Apoptosis Detection Kit (Biolegend) according to the manufacturer’s protocol. Antibodies are detailed in Supplementary Table S5. Data were acquired on a LSRII flow cytometer (BD) or a LSRFortessa (BD) and analyzed with FlowJo software V10. Chemokines from serum, urine, bladder supernatant and tumor cell supernatant of mice were analyzed using LEGENDplex^TM^ kits following manufacturer’s recommendations (#740683 and #740451; Biolegend). CXCL12 from serum and urine of patients and healthy donors were analyzed using LEGENDplex^TM^ kits following manufacturer’s recommendations 741170; Biolegend) and acquired using a CytoFlex (BD) instrument and data were analyzed with the LEGENDplex™ Data Analysis software (v. 8.0). For macrophage isolation, extracellular markers were stained for 30 min in homemade SORT buffer. DAPI was added to exclude dead cells just before running samples. CD45^+^CD11b^+^CD4^-^CD8^-^F4/80^+^MHCII^low^ cells and CD45^+^CD11b^+^CD4^-^CD8^-^F4/80^+^MHCII^high^ cells (Figure S1) were then isolated by sorting on MoFlo Astrios EQ (Beckman coulter) at the Flow Cytometry Facility of the University of Lausanne. Cells were collected in RNA later buffer (#AM7020, Invitrogen) before RNA extraction as described above.

### Immunohistochemistry of mouse sections

Murine tumors were freshly frozen in OCT, and cryostat sections (8 μm thick) were fixed in ice-cold acetone for 10 minutes before rehydration in PBS. Sections were blocked in a buffer consisting of 0.1% (wt/vol) bovine serum albumin (BSA), 1% (vol/vol) mouse serum (M5905, Sigma), and 1% (vol/vol) normal donkey serum (D9663, Sigma) for 30 minutes at room temperature (RT). Immunostaining was conducted using the primary and secondary antibodies detailed in Supplementary Table S5. Antibody dilutions were prepared in the blocking buffer. Images were acquired using a NanoZoomer S60 slide scanner. Images were then analyzed with QuPath software (54).

### In situ RNA hybridization and immunofluorescence microscopy of mice sections

RNA-Scope was performed using the RNAscope Multiplex Fluorescent Detection Kit v2 kit (323110, ACD) and RNAscope H_2_O_2_ and protease Reagents kit (322381, ACD) according to the manufacturer’s instructions. Briefly, tissue sections were rehydrated in PBS 1X for 5 min at RT, incubated 30 min at 60°C in a HybEZ II oven, and fixed for 15 min at 4°C in 4% PFA. Tissue sections were treated with H_2_O_2_ for 10 min at RT followed by incubation in target retrieval reagents solution for 11 min at 90°C and protease III solution for 30 min at 40°C. Then, sections were incubated with the RNAscope™ Probe-Mm-Cxcl12-C2 (ref 422711-C2 ACDBio), positive probe (RNAscope 3-plex positive control probe-Mm) and negative probe (RNAscope 3-plex negative control Probe). The hybridization procedure was performed for 2 hours at 40°C. Sequential amplification steps were performed according to manufacturer’s instructions using Amp1, Amp2 and Amp3 solutions at 40°C. Last, tissue sections were incubated with Opal650 (OP-001005 Akoya Biosciences) for 30 min at 40°C. Then, sections were incubated with Hoechst for nuclear staining. Autofluorescence was removed with the True Black Kit (92401 TrueBlack Lipofuscin Autofluorescence Quencher) according to the manufacturer’s instructions. Mounting was performed using the mounting medium ProLong Gold antifade reagent (P36934 Invitrogen). Images were acquired using a LSM780 confocal microscope and processed using the open-source digital image analysis software QuPath v0.2.3 (54).

### Immunohistochemistry analyses of human bladder sections

BC patients’ tissue sections were obtained from the Biobank of Institute of Pathology at the CHUV in the frame of the study protocol n°2019/00882 approved by Commission cantonale d’éthique de la recherche sur l’être humain, canton of Vaud (CER-VD). Tumor areas were defined by the pathologist based on histology sections. Patients’ clinical information is summarized in Supplementary Table S4.

Formalin-Fixed Paraffin-Embedded (FFPE) human tissue sections were subjected to heat-induced antigen retrieval (HIER) using a citrate buffer at pH 6.0. Sections were blocked, and antibody dilutions were prepared in a buffer containing 0.1% BSA (wt/vol), 1% (vol/vol) human serum (S1, Sigma), and 1% (vol/vol) normal donkey serum. At the end of the staining, sections were quenched with TrueVIEW® Autofluorescence Quenching Kit according to manufacturer’s instructions. Antibodies are detailed in Supplementary Table S5. Images were acquired using a NanoZoomer S60 slide scanner. Images were then analyzed with QuPath software (54). Regions of interest were drawn, and cells were defined by the “cell detection” parameter on the DAPI as the detection channel, with 12µm of background radius, 2.05 µm of Sigma, 900 threshold and 2µm cell expansion. CD68 was qualified positive for a measurement of Cytoplasm with a mean threshold of 1400. CD204 was qualified positive for a measurement of Cytoplasm with a mean threshold of 289. CXCR4 was qualified positive for a measurement of Cytoplasm with a mean threshold of 1050. CD8 was qualified positive for a measurement of Cytoplasm with a mean threshold of 4,1. CD3 was qualified positive for a measurement of Cytoplasm with a mean threshold of 15,38.

### RNA sequencing

Total RNA from whole bladder and sorted macrophages were extracted using the RNeasy Plus Micro Kit (#74034, Qiagen) according to manufacturer’s instructions. RNA quality was assessed using Fragment Analyzer System (Agilent) and RNA Kit (#DNF-471, Agilent). RNA samples were polyA-enriched and libraries were prepared using the Illumina TruSeq® Stranded RNA kit. Single-end (125 bp) RNA sequencing with a depth of approximately 20–30 million reads per sample was performed on Illumina’s Hi-Seq 2500 platform at the Genomic Technologies Facility of Lausanne.

### Bioinformatics analysis

RNA-seq quantification was performed using kallisto (55). In brief, target transcript sequences were obtained from ENSEMBLE (GRCm38.p6), and the abundances of transcripts were quantified using kallisto 0.44.0 with sequence-based bias correction. All other parameters were set to default when running kallisto. Kallisto’s transcript-level estimates were further summarized at the gene-level using tximport 1.8.0 from Bioconductor (56). Lowly abundant genes were filtered out prior to downstream analyses. For tumor cell lines, unwanted variation was estimated using the SVA 3.30.0 package from Bioconductor (57). The number of factors of unwanted variation to be estimated from the data was set to 2. Multidimensional scaling plot was generated using the limma package from Bioconductor (58), with top 500 variable genes chosen separately for each pairwise comparison. Differential expression analysis was performed using DESeq2 1.22.0 from Bioconductor (59). Surrogate variables of unwanted variation were included as additional covariates in the design formula when analyzing tumor cell lines. Significant genes were identified using FDR<0.05. Single-sample gene set enrichment analysis [ssGSEA, (60)] was performed using the GSVA 1.30.0 package from Bioconductor (61), with regularized log-transformed (rlog) normalized data obtained from DESeq2.

### Database analysis

TCGA database analyses were performed on the GEPIA2 website (62). Pro-tumor TAM signature was taken from Cassetta and colleagues (31).

### Graphics and Statistics

GraphPad Prism 10 software was used to generate graph and to perform statistical analyses. Used tests are specified in the legend of each Figure. For non-significant differences, p values are absent, while statistically significant results, p values are added on the figures: *p<0.05; **p<0.01; ***p<0.001; ****p<0.0001. Heatmaps were generated with the Morpheus software (https://software.broadinstitute.org/morpheus) or GraphPad Prism 10 software.

## Supplementary figure legends

**Figure S1.** Identification of mouse and human macrophages. (**A**) Gating strategy for mouse macrophages sorting. MHCII^low^ macrophages were gated on CD45^+^CD11b^+^CD3^-^F4/80^+^MHCII^low^ cells and MHCII^high^ macrophages were gated on CD45^+^CD11b^+^CD3^-^F4/80^+^MHCII^high^ cells. (**B**) Dot plot representing the ssGSEA enrichment score of the sorted MHCII^high^ and MHCII^low^ macrophages between healthy bladders and bladders with NMIBC, MIBC and advanced MIBC. Each dot represents an individual mouse. One-way ANOVA, followed by Tukey’s HSD test. (**C**) Human macrophages were gated on DUMP^-^ CD45^+^CD11b^+^CD11c^+^CD14^+^HLA-DR^high^ cells.

**Figure S2.** Ligand-receptor expressions in BC-bearing mice. (**A**) Representative flow cytometry contour plots and quantification of CSF1R and CCR2 on MHCII^low^ and MHCII^high^ macrophages in bladder of healthy, NMIBC-, MIBC- and advanced MIBC-bearing mice. (**B**) Representative flow cytometry contour plot of Ly6C over Ly6G expressions on myeloid cells and NK1.1 over CD3 expressions on immune cells and the related histograms of CXCR4 expression on the different populations in bladder of MIBC-bearing mice. (**C**) Bar plot of CXCR4 geometric mean fluorescent intensity (GMFI) on CD8^+^, NK^+^, B220^+^, FoxP3^+^CD4^+^, FoxP3^-^CD4^+^, Ly6G^+^ and Ly6C^high^F480^-^ cell populations in blood of MIBC-bearing mice. (**D**) Bar plot of CXCR4^+^ expression on BMDMs in control (CT) conditions or after the addition of tumor conditioned media (TCM) from advanced MIBC-bearing mice. Each dot represents an individual experiment, and bars represent the mean ± SD. Unpaired t-tests.

**Figure S3.** Blocking CSF1-CSF1R and CCL2-CCR2 pathways in MIBC-bearing mice. (**A**) Timeline of anti-CSF1R or anti-CCL2/CCR2 treatments in MIBC-bearing mice. (**B**) Kaplan-Meier curve of control (mock) and anti-CSF1R or anti-CCL2/CCR2 treated mice. n=7. Log-rank test.

## Notes

### Competing Interest Statement

The authors have declared no competing interest.

