## Supplementary material for "Targeting CXCR4-expressing TAMs in muscle-invasive bladder cancer to enhance tumor control after immunotherapy": Suppl. Figure 1 to 3; Suppl. tables 1 to 5

Figure S1

A

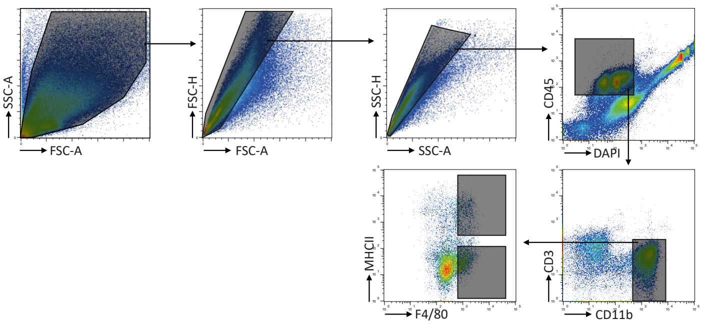


B

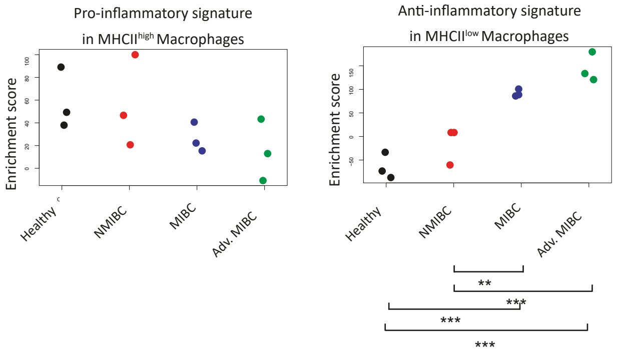


C

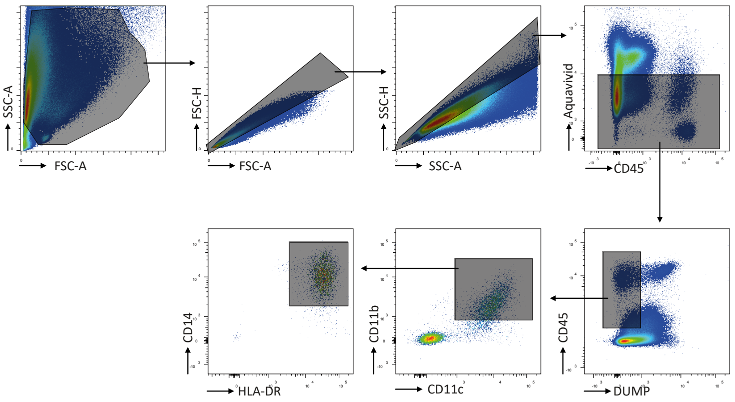

Figure S2

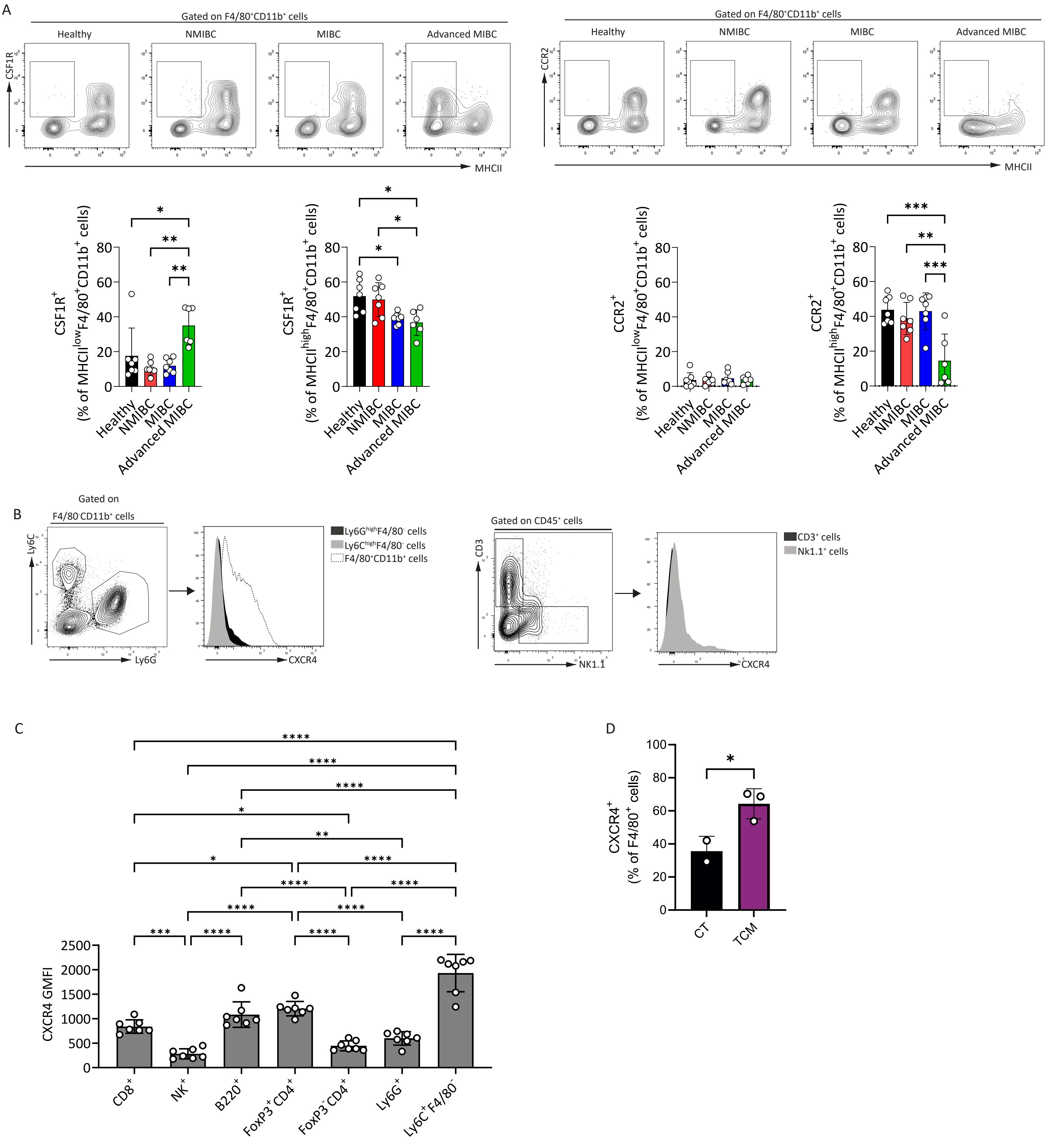
A
Gated on F4/80+CD11b+ cells
Healthy
NMIBC
MIBC
Advanced MIBC
CSF1R
MHCII
CCR2
CSF1R+ (% of MHCIIlowF4/80+CD11b+ cells)
CSF1R+ (% of MHCIIhighF4/80+CD11b+ cells)
CCR2+ (% of MHCIIlowF4/80+CD11b+ cells)
CCR2+ (% of MHCIIhighF4/80+CD11b+ cells)
B
Gated on F4/80-CD11b+ cells
Ly6C
Ly6G
Ly6GhighF4/80- cells
Ly6ChighF4/80- cells
F4/80+CD11b+ cells
CXCR4
Gated on CD45+ cells
CD3
NK1.1
CD3+ cells
Nk1.1+ cells
C
CXCR4 GMFI
CD8+
NK+
B220+
FoxP3+ CD4+
FoxP3- CD4+
Ly6G+
Ly6C+ F4/80-
D
CXCR4+ (% of F4/80+ cells)
CT
TCM

Figure S3

A

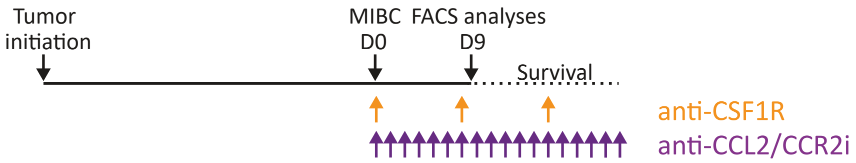
Tumor
initiation
MIBC
D0
FACS analyses
D9
Survival
anti-CSF1R
anti-CCL2/CCR2i

B

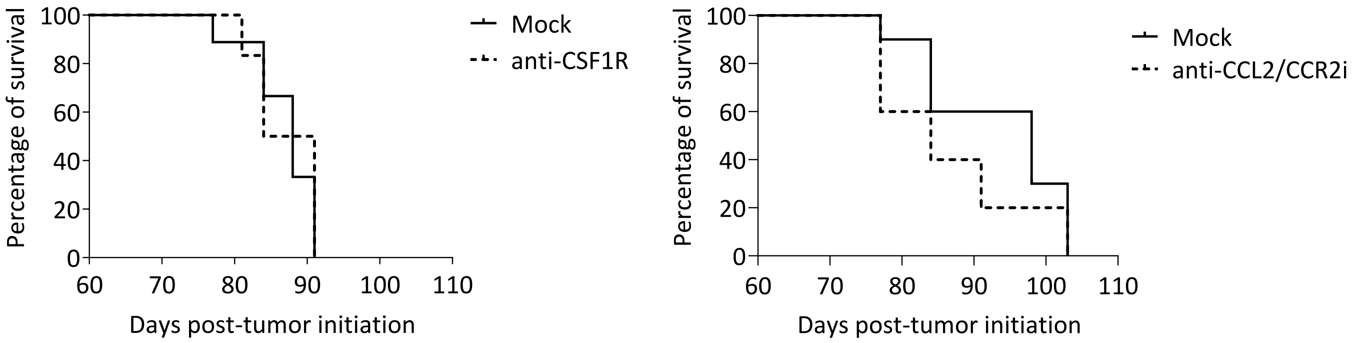
Percentage of survival
100
80
60
40
20
0
60
70
80
90
100
110
Days post-tumor initiation
Mock
anti-CSF1R
Mock
anti-CCL2/CCR2i

### Supplementary tables

#### Supplemental Table S1. Chemokines concentrations analyzed by Legend Plex

Bladder supernatant

| | CT | CT | CT | CT | CT | CT | 5w | 5w | 5w | 5w | 5w | 5w | 9w | 9w | 9w | 9w | 9w | 9w | 13w | 13w | 13w | 13w | 13w | 13w |
|---|---|---|---|---|---|---|---|---|---|---|---|---|---|---|---|---|---|---|---|---|---|---|---|---|
| CCL5 | 28 | 33 | 26 | 226 | 28 | 45 | 121 | 27 | 92 | 39 | 23 | 100 | 185 | 40 | 76 | 80 | 100 | 64 | 222 | 46 | 42 | 40 | 77 | 17 |
| CCL20 | 86 | 26 | 36 | 36 | 26 | 31 | 16 | 36 | 64 | 55 | 31 | 20 | 79 | 68 | 46 | 36 | 31 | 71 | 16 | 51 | 46 | 26 | 31 | 16 |
| CCL11 | 613 | 283 | 571 | 1664 | 508 | 767 | 571 | 155 | 773 | 314 | 453 | 348 | 2176 | 1443 | 1066 | 1176 | 613 | 594 | 1085 | 2039 | 986 | 989 | 685 | 453 |
| CCL17 | 250 | 93 | 157 | 165 | 103 | 93 | 42 | 139 | 182 | 121 | 42 | 73 | 213 | 121 | 103 | 83 | 112 | 112 | 103 | 157 | 121 | 157 | 83 | 112 |
| CXCL1 | 5416 | 5713 | 18499 | 2285 | 6980 | 7737 | 7170 | 18032 | 8698 | 11002 | 14021 | 2258 | 22858 | 22858 | 22858 | 13937 | 22858 | 21756 | 14764 | 22858 | 21374 | 22858 | 15002 | 13137 |
| CCL2 | 5114 | 6486 | 10101 | 23917 | 5164 | 8245 | 3376 | 5363 | 3472 | 4400 | 5065 | 9639 | 15252 | 13197 | 9157 | 9319 | 14620 | 6218 | 8539 | 7299 | 6026 | 9066 | 6716 | 3018 |
| CXCL9 | 78 | 78 | 78 | 145 | 78 | 106 | 179 | 78 | 166 | 78 | 78 | 261 | 297 | 128 | 91 | 78 | 234 | 78 | 78 | 78 | 78 | 124 | 78 | 78 |
| CXCL10 | 2823 | 1951 | 1027 | 4758 | 1384 | 2188 | 1193 | 443 | 1033 | 960 | 1088 | 768 | 2946 | 1512 | 1530 | 1587 | 1806 | 1407 | 1658 | 1097 | 1519 | 2029 | 1513 | 396 |
| CCL3 | 148 | 125 | 85 | 227 | 78 | 173 | 56 | 80 | 75 | 105 | 83 | 190 | 196 | 181 | 282 | 146 | 120 | 76 | 218 | 319 | 105 | 153 | 105 | 26 |
| CCL4 | 16 | 14 | 13 | 66 | 13 | 35 | 13 | 15 | 22 | 13 | 13 | 45 | 63 | 40 | 64 | 26 | 26 | 13 | 39 | 68 | 27 | 41 | 16 | 13 |
| CXCL13 | 495 | 415 | 599 | 1157 | 989 | 1455 | 698 | 114 | 1137 | 520 | 603 | 979 | 3038 | 1317 | 2011 | 1339 | 1123 | 199 | 574 | 1141 | 676 | 1102 | 263 | 394 |
| CXCL5 | 125 | 79 | 575 | 979 | 328 | 562 | 198 | 500 | 136 | 260 | 249 | 310 | 5254 | 2434 | 3477 | 1223 | 1397 | 1090 | 1707 | 6050 | 1610 | 1707 | 1034 | 963 |
| CCL22 | 1249 | 473 | 522 | 1989 | 1381 | 1130 | 636 | 120 | 802 | 417 | 338 | 702 | 4545 | 1025 | 1272 | 581 | 830 | 664 | 490 | 2093 | 448 | 577 | 469 | 767 |
| CSF1 | 11803 | 16690 | 15541 | 24240 | 16260 | 18102 | 10006 | 10814 | 4356 | 11764 | 14661 | 13267 | 9931 | 24240 | 6446 | 14280 | 24240 | 10196 | 22205 | 24240 | 24240 | 22590 | 20283 | 8625 |
| CXCL12 | 38 | 46 | 38 | 31 | 38 | 29 | 38 | 29 | 21 | 46 | 13 | 4 | 109 | 64 | 117 | 56 | 82 | 64 | 59 | 162 | 64 | 144 | 109 | 38 |

Serum

| | CT | CT | CT | CT | CT | CT | CT | CT | CT | CT | 5w | 5w | 5w | 5w | 5w | 5w | 5w | 5w | 5w | 5w |
|---|---|---|---|---|---|---|---|---|---|---|---|---|---|---|---|---|---|---|---|---|
| CCL5 | 15 | 12 | 0 | 0 | 12 | 14 | 0 | 13 | 11 | 0 | 9 | 15 | 11 | 14 | 26 | 10 | 11 | 0 | 0 | 9 |
| CCL20 | 51 | 0 | 12 | 22 | 0 | 17 | 9 | 0 | 0 | 0 | 0 | 12 | 0 | 0 | 0 | 7 | 0 | 17 | 0 | 9 |
| CCL11 | 4476 | 3185 | 2945 | 1766 | 3960 | 969 | 667 | 1746 | 3193 | 1982 | 2611 | 6518 | 4184 | 9514 | 2742 | 1753 | 1815 | 2036 | 1685 | 1821 |
| CCL17 | 16 | 0 | 41 | 16 | 0 | 41 | 0 | 36 | 0 | 0 | 0 | 12 | 71 | 0 | 84 | 0 | 47 | 25 | 47 | 0 |
| CXCL1 | 214 | 68 | 130 | 180 | 103 | 92 | 59 | 100 | 81 | 62 | 89 | 160 | 137 | 0 | 130 | 47 | 87 | 51 | 31 | 93 |
| CCL2 | 24 | 31 | 0 | 31 | 31 | 0 | 31 | 28 | 0 | 11 | 0 | 21 | 22 | 0 | 34 | 22 | 0 | 28 | 17 | 24 |
| CXCL9 | 102 | 8 | 0 | 28 | 0 | 28 | 0 | 0 | 0 | 61 | 0 | 98 | 16 | 13 | 8 | 0 | 11 | 0 | 0 | 13 |
| CXCL10 | 8 | 0 | 0 | 0 | 0 | 0 | 0 | 0 | 0 | 0 | 0 | 5 | 0 | 0 | 0 | 0 | 0 | 0 | 0 | 0 |
| CCL3 | 0 | 0 | 0 | 0 | 0 | 0 | 0 | 0 | 0 | 0 | 0 | 3 | 0 | 0 | 0 | 0 | 0 | 0 | 0 | 0 |
| CCL4 | 0 | 0 | 0 | 0 | 0 | 0 | 4 | 0 | 0 | 0 | 0 | 16 | 0 | 0 | 0 | 0 | 0 | 0 | 0 | 0 |
| CXCL13 | 1294 | 643 | 1094 | 778 | 958 | 609 | 437 | 990 | 510 | 691 | 945 | 1278 | 1022 | 996 | 752 | 3442 | 644 | 974 | 828 | 515 |
| CXCL5 | 72 | 424 | 216 | 414 | 153 | 143 | 486 | 430 | 75 | 161 | 82 | 752 | 53 | 27 | 233 | 319 | 415 | 546 | 244 | 170 |
| CCL22 | 290 | 347 | 55 | 44 | 227 | 26 | 0 | 84 | 319 | 201 | 268 | 468 | 377 | 1039 | 170 | 29 | 97 | 119 | 58 | 47 |
| CSF1 | 10455 | 7150 | 14250 | 14159 | 12350 | 15787 | 9583 | 16682 | 16163 | 18388 | 11606 | 7357 | 9757 | 17272 | 11664 | 13541 | 13141 | 8764 | 12546 | 16570 |
| CXCL12 | 140 | 170 | 197 | 162 | 187 | 149 | 196 | 120 | 204 | 110 | 90 | 140 | 122 | 144 | 170 | 94 | 149 | 136 | 140 | 166 |

| | 9w | 9w | 9w | 9w | 9w | 9w | 9w | 9w | 9w | 9w | 13w | 13w | 13w | 13w | 13w | 13w | 13w | 13w | 13w | 13w |
|---|---|---|---|---|---|---|---|---|---|---|---|---|---|---|---|---|---|---|---|---|
| CCL5 | 15 | 15 | 9 | 11 | 84 | 6 | 0 | 0 | 0 | 0 | 50 | 24 | 34 | 11 | 14 | 27 | 10 | 20 | 14 | 18 |
| CCL20 | 0 | 0 | 0 | 0 | 9 | 0 | 0 | 0 | 7 | 0 | 46 | 6 | 0 | 6 | 12 | 0 | 0 | 6 | 0 | 0 |
| CCL11 | 4428 | 1748 | 3122 | 3166 | 1595 | 4176 | 2062 | 2423 | 4564 | 1598 | 2017 | 2261 | 3389 | 2455 | 2165 | 2508 | 4482 | 2002 | 1140 | 1544 |
| CCL17 | 84 | 0 | 36 | 0 | 0 | 0 | 0 | 78 | 0 | 0 | 30 | 0 | 78 | 0 | 30 | 0 | 47 | 12 | 25 | 59 |
| CXCL1 | 77 | 90 | 180 | 84 | 85 | 11 | 17 | 128 | 82 | 325 | 189 | 134 | 170 | 0 | 256 | 166 | 521 | 278 | 148 | 323 |
| CCL2 | 17 | 48 | 28 | 0 | 32 | 42 | 11 | 34 | 44 | 19 | 17 | 10 | 211 | 29 | 36 | 42 | 34 | 48 | 42 | 78 |
| CXCL9 | 0 | 0 | 0 | 65 | 11 | 38 | 25 | 28 | 228 | 91 | 22 | 25 | 0 | 0 | 0 | 87 | 0 | 0 | 16 | 16 |
| CXCL10 | 0 | 0 | 0 | 4 | 0 | 0 | 0 | 0 | 0 | 5 | 175 | 0 | 333 | 0 | 0 | 427 | 0 | 50 | 0 | 79 |
| CCL3 | 0 | 0 | 0 | 0 | 0 | 0 | 0 | 0 | 0 | 0 | 0 | 0 | 0 | 0 | 0 | 0 | 0 | 0 | 0 | 0 |
| CCL4 | 0 | 0 | 6 | 14 | 229 | 17 | 0 | 6 | 0 | 0 | 74 | 13 | 9 | 0 | 0 | 25 | 0 | 8 | 0 | 31 |
| CXCL13 | 1156 | 215 | 686 | 9945 | 1104 | 5949 | 1324 | 889 | 863 | 724 | 2636 | 201 | 8430 | 3016 | 1060 | 10785 | 1385 | 5556 | 3802 | 3109 |
| CXCL5 | 45 | 0 | 368 | 466 | 260 | 24 | 179 | 329 | 110 | 113 | 222 | 86 | 85 | 214 | 333 | 190 | 237 | 192 | 98 | 135 |
| CCL22 | 409 | 79 | 278 | 223 | 55 | 50 | 172 | 167 | 343 | 39 | 221 | 269 | 92 | 112 | 146 | 160 | 487 | 361 | 94 | 60 |
| CSF1 | 12421 | 22888 | 15495 | 12730 | 15674 | 10187 | 4448 | 12232 | 13114 | 21475 | 16227 | 6102 | 16202 | 12289 | 19258 | 19290 | 17988 | 22687 | 16762 | 22888 |
| CXCL12 | 66 | 127 | 200 | 153 | 94 | 80 | 90 | 170 | 170 | 94 | 359 | 122 | 136 | 94 | 489 | 232 | 232 | 236 | 140 | 305 |

Tumor cell supernatant

| | 5w | 5w | 5w | 5w | 5w | 9w | 9w | 9w | 9w | 9w | 13w | 13w | 13w | 13w | 13w |
|---|---|---|---|---|---|---|---|---|---|---|---|---|---|---|---|
| CCL5 | 0 | 0 | 8 | 0 | 0 | 0 | 6 | 15 | 9 | 8 | 5 | 0 | 5 | 0 | 0 |
| CCL20 | 31 | 0 | 29 | 16 | 33 | 33 | 35 | 37 | 0 | 22 | 44 | 41 | 19 | 22 | 37 |
| CCL11 | 0 | 0 | 0 | 0 | 0 | 7 | 0 | 18 | 7 | 17 | 9 | 6 | 9 | 18 | 9 |
| CCL17 | 0 | 0 | 0 | 0 | 0 | 0 | 0 | 0 | 0 | 0 | 0 | 0 | 0 | 0 | 0 |
| CXCL1 | 90 | 20 | 116 | 61 | 78 | 152 | 228 | 104 | 130 | 58 | 279 | 57 | 849 | 76 | 68 |
| CCL2 | 43 | 26 | 86 | 51 | 57 | 96 | 192 | 105 | 177 | 58 | 111 | 49 | 706 | 78 | 113 |
| CXCL9 | 22 | 0 | 19 | 0 | 33 | 0 | 0 | 0 | 0 | 0 | 0 | 0 | 0 | 0 | 0 |
| CXCL10 | 9 | 5 | 12 | 0 | 0 | 42 | 35 | 29 | 69 | 12 | 37 | 0 | 17 | 0 | 13 |
| CCL3 | 0 | 0 | 0 | 0 | 0 | 0 | 0 | 0 | 0 | 0 | 0 | 0 | 4 | 0 | 2 |
| CCL4 | 6 | 0 | 4 | 0 | 4 | 0 | 6 | 0 | 0 | 0 | 0 | 5 | 43 | 0 | 7 |
| CXCL13 | 0 | 0 | 9 | 0 | 9 | 0 | 0 | 0 | 8 | 0 | 0 | 0 | 9 | 0 | 0 |
| CXCL5 | 23 | 10 | 10 | 0 | 22 | 0 | 10 | 18 | 11 | 10 | 11 | 0 | 16 | 10 | 15 |
| CCL22 | 14 | 8 | 11 | 0 | 11 | 0 | 8 | 8 | 11 | 23 | 12 | 0 | 17 | 9 | 11 |
| CSF1 | 5683 | 9748 | 10054 | 8587 | 9704 | 8423 | 12007 | 9862 | 8894 | 8046 | 2768 | 5797 | 6823 | 6863 | 6388 |
| CXCL12 | 188 | 8 | 188 | 336 | 336 | 510 | 982 | 510 | 510 | 72 | 226 | 238 | 888 | 348 | 576 |

Urine

| | CT | CT | CT | CT | CT | CT | CT | 5w | 5w | 5w | 5w | 5w | 5w | 5w | 9w | 9w | 9w | 9w | 9w | 9w | 9w | 13w | 13w | 13w | 13w | 13w | 13w |
|---|---|---|---|---|---|---|---|---|---|---|---|---|---|---|---|---|---|---|---|---|---|---|---|---|---|---|---|
| CCL5 | 10 | 3 | 6 | 3 | 7 | 3 | 3 | 3 | 3 | 3 | 3 | 3 | 3 | 4 | 5 | 3 | 7 | 3 | 3 | 3 | 5 | 3 | 4 | 3 | 3 | 3 | 3 |
| CCL20 | 4 | 7 | 3 | 3 | 10 | 3 | 3 | 11 | 3 | 3 | 3 | 3 | 3 | 3 | 1 | 1 | 1 | 1 | 1 | 1 | 1 | 3 | 3 | 4 | 5 | 3 | 3 |
| CCL11 | 3 | 3 | 14 | 3 | 3 | 3 | 12 | 3 | 3 | 3 | 3 | 3 | 3 | 3 | 4 | 8 | 3 | 3 | 3 | 3 | 5 | 8 | 3 | 7 | 3 | 16 | 3 |
| CCL17 | 20 | 20 | 20 | 34 | 34 | 28 | 20 | 20 | 20 | 20 | 20 | 23 | 22 | 20 | 20 | 20 | 20 | 20 | 20 | 20 | 20 | 20 | 20 | 20 | 20 | 20 | 20 |
| CXCL1 | 8 | 8 | 50 | 8 | 8 | 8 | 8 | 4 | 4 | 4 | 4 | 4 | 4 | 5 | 4 | 4 | 25 | 4 | 4 | 4 | 4 | 4 | 4 | 4 | 4 | 4 | 4 |
| CCL2 | 14 | 5 | 5 | 5 | 5 | 7 | 5 | 5 | 5 | 5 | 5 | 6 | 7 | 5 | 5 | 5 | 5 | 5 | 5 | 5 | 5 | 7 | 5 | 5 | 5 | 5 | 5 |
| CXCL9 | 3 | 4 | 22 | 6 | 3 | 5 | 3 | 17 | 7 | 3 | 3 | 3 | 3 | 3 | 3 | 15 | 3 | 3 | 3 | 4 | 4 | 6 | 22 | 9 | 3 | 4 | 3 |
| CXCL10 | 18 | 7 | 8 | 7 | 4 | 4 | 5 | 41 | 14 | 16 | 12 | 19 | 28 | 23 | 11 | 40 | 23 | 17 | 64 | 10 | 32 | 35 | 28 | 43 | 10 | 169 | 43 |
| CCL3 | 3 | 2 | 2 | 3 | 1 | 2 | 1 | 1 | 1 | 1 | 1 | 3 | 1 | 1 | 1 | 1 | 1 | 1 | 1 | 1 | 2 | 2 | 1 | 2 | 2 | 1 | 5 |
| CCL4 | 132 | 2 | 4 | 2 | 2 | 38 | 2 | 2 | 2 | 2 | 2 | 109 | 2 | 13 | 2 | 6 | 2 | 2 | 2 | 23 | 6 | 2 | 2 | 2 | 2 | 2 | 2 |
| CXCL13 | 3 | 1 | 3 | 1 | 1 | 1 | 1 | 1 | 1 | 1 | 1 | 3 | 2 | 1 | 1 | 2 | 1 | 1 | 2 | 1 | 3 | 3 | 1 | 2 | 1 | 10 | 1 |
| CXCL5 | 8 | 27 | 32 | 39 | 35 | 6 | 6 | 23 | 6 | 26 | 6 | 18 | 18 | 6 | 7 | 9 | 17 | 4 | 11 | 26 | 41 | 22 | 26 | 72 | 69 | 21 | 27 |
| CCL22 | 4 | 4 | 4 | 7 | 4 | 4 | 5 | 4 | 7 | 4 | 4 | 4 | 4 | 4 | 4 | 4 | 4 | 4 | 4 | 4 | 4 | 4 | 4 | 5 | 4 | 4 | 4 |
| CSF1 | 1347 | 1936 | 711 | 3010 | 2478 | 2441 | 1494 | 4243 | 557 | 349 | 1051 | 708 | 1077 | 747 | 488 | 2003 | 747 | 821 | 667 | 484 | 1075 | 455 | 465 | 441 | 691 | 347 | 901 |
| CXCL12 | 48 | 50 | 55 | 31 | 56 | 43 | 48 | 31 | 34 | 30 | 32 | 29 | 59 | 45 | 66 | 55 | 45 | 58 | 49 | 30 | 55 | 90 | 66 | 65 | 79 | 79 | 81 |

**Supplemental Table S2. Patients' clinical information from biopsies and blood samples.** Here the pathological (p) stage was determined according to the AJCC staging system used for bladder cancer, with T (primary tumor), N (lymph node(s)) and M (metastasis). pT1: the cancer has grown into the layer of connective tissue under the lining layer of the bladder, but it has not reached the layer of muscle in the bladder wall. Cancer grade describes how abnormal the bladder cancer cells look under a microscope and how quickly the cancer cells are likely to grow and spread.

| Parameter | Total cases | Percentage % |
|---|---|---|
| Sex | | |
| Female | 22 | 39,29 |
| Male | 34 | 60,71 |
| Age | | |
| <50 yo | 2 | 3,57 |
| 50-70 | 20 | 35,71 |
| >70 | 34 | 60,71 |
| Tumor stage | | |
| ≤ pT1 | 39 | 69,64 |
| >pT1 | 17 | 30,36 |
| Tumor grade | | |
| Low grade | 21 | 37,50 |
| High grade | 35 | 62,50 |

**Supplemental Table S3. Patients' clinical information from urine samples.** Here the pathological (p) stage was determined according to the AJCC staging system used for bladder cancer, with T (primary tumor), N (lymph node(s)) and M (metastasis). pT1: the cancer has grown into the layer of connective tissue under the lining layer of the bladder, but it has not reached the layer of muscle in the bladder wall. Cancer grade describes how abnormal the bladder cancer cells look under a microscope and how quickly the cancer cells are likely to grow and spread.

| Parameter | Total cases | Percentage % |
|---|---|---|
| Sex | | |
| Female | 2 | 9,09 |
| Male | 20 | 90,91 |
| Age | | |
| <50 yo | 1 | 4,55 |
| 50-70 | 17 | 77,27 |
| >70 | 4 | 18,18 |
| Tumor stage | | |
| ≤ pT1 | 21 | 95,45 |
| >pT1 | 1 | 4,55 |
| Tumor grade | | |
| Low grade | 11 | 50,00 |
| High grade | 11 | 50,00 |

**Supplemental Table S4. Patients' clinical information from paraffin-embedded tissue sections.** Here the pathological (p) stage was determined according to the AJCC staging system used for bladder cancer, with T (primary tumor), N (lymph node(s)) and M (metastasis). pT2: the cancer has grown into the inner or outer muscle layer of the bladder wall. pT3: the cancer has grown through the muscle layer of the bladder and into the layer of fatty tissue that surrounds the bladder. pT4: the cancer has at least grown into the layer of connective tissue under the lining of the bladder wall. pNx: pathologists were unable to find out if bladder cancer has spread to the lymph nodes. pN0: no lymph nodes containing cancer cells. pN1: cancer cells infitrated into 1 lymph node in the true pelvis. pN2: cancer cells infiltrated into 2 or more lymph nodes in the true pelvis. Cancer grade describes how abnormal the bladder cancer cells look under a microscope and how quickly the cancer cells are likely to grow and spread.

| Parameter | Total cases | Percentage % |
|---|---|---|
| Sex | | |
| Female | 3 | 12,00 |
| Male | 22 | 88,00 |
| Tumor stage | | |
| pT2 | 7 | 28,00 |
| pT3 | 14 | 56,00 |
| pT4 | 4 | 16,00 |
| Lymph node metastasis | | |
| pNx | 4 | 16,00 |
| pN0 | 13 | 52,00 |
| pN1 | 4 | 16,00 |
| pN2 | 4 | 16,00 |
| Tumor grade | | |
| Unknown | 4 | 16,00 |
| Low grade | 0 | 0,00 |
| High grade | 21 | 84,00 |

**Supplemental Table S5. Antibodies used for mouse and human cytometry and immunohistochemistry**

| Marker | Fluo | Dilution | Reference |
|---|---|---|---|
| **Mouse flow cytometry Extracellular** | | | |
| CD4 | APC-Cy7 | 1:300 | 47-0042-82 |
| CD4 | BUV496 | 1:500 | 364-0042-82 |
| CD45.2 | BV650 | 1:100 | 109847 |
| CD45.2 | BUV805 | 1:50 | 368-0454-82 |
| CD8 | BV786 | 1:200 | 100750 |
| F4/80 | PE TexasRed | 1:1000 | 61-4801-82 |
| F4/80 | APC-Cy7 | 1:100 | 123118 |
| Ly6C | PercpCy5.5 | 1:200 | 45-5932-82 |
| Ly6G | BV650 | 1:200 | 127641 |
| Ly6G | AlFl700 | 1:200 | 127621 |

| | | | |
|---|---|---|---|
| MHCII | BV785 | 1:1000 | 107645 |
| PDL1 | BV711 | 1:200 | 124319 |
| CD11b | APC-Cy7 | 1:200 | 47-0112-82 |
| CD11b | BV605 | 1:500 | 101257 |
| CD11b | PE Cy5 | 1:1000 | 15-0112-83 |
| CD25 | PE | 1:2000 | 12-0251-82 |
| B220 | BV570 | 1:50 | 103237 |
| NK1.1 | APC | 1:100 | 108710 |
| CD3 | eFl506 | 1:50 | 69-0032-82 |
| CD204 | PE Cy7 | 1:200 | 25-2046-80 |
| CXCR4 | PE | 1:50 | FAB21651P |
| CXCR4 | BV711 | 1:100 | 146505 |

| **Mouse flow cytometry Intracellular** | | | |
|---|---|---|---|
| Arg1 | APC | 1:50 | IC5868A |
| CD206 | APC | 1:100 | 141712 |
| FoxP3 | BV421 | 1:200 | 48-5773-82 |
| Ki67 | BV421 | 1:100 | 652411 |

| **Human flow cytometry Extracellular** | | | |
|---|---|---|---|
| CXCR4 | BV785 | 1:100 | 306529 |
| CD163 | BV711 | 1:50 | 333630 |
| CD169 | BV605 | 1:100 | 346009 |
| CD14 | BV421/PB | 1:200 | 367121 |
| Dump CD3 | PerCPCy5.5 | 1:200 | 317335 |
| CD19 | | 1:50 | 302230 |
| CD56 | | 1:50 | 318322 |
| CD66b | | 1:200 | 396913 |
| HLA-DR | FITC | 1:250 | 327006 |
| CD11c | PeCy7 | 1:200 | 371507 |
| CD16 | ECD | 1:300 | 302053 |
| CD45 | APC-Cy7 | 1:100 | 368515 |
| CD11b | A700 | 1:50 | 301356 |
| Tie-2 | APC | 1:50 | 334209 |

| **Mouse IHC** | | | |
|---|---|---|---|
| CXCR4 | | 1:500 | ab181020 |
| F4/80 | | 1:4 | Homemade |
| CD3 | | 1 :300 | ab11089 |
| CD8b | | 1 :100 | 53-0083-82 |
| SDF1 | | 1 :100 | ab155090 |
| anti-rabbit IgG | | 1 :500 | A32794 |
| anti-goat IgG | | 1:600 | 705-545-003 |

| | | | |
|---|---|---|---|
| anti-rat IgG | | 1 :500 | 712-136-153 |

| **Human IHC** | | | |
|---|---|---|---|
| CD68 | | 1:50 | ab213363 |
| CXCR4 | | 1:100 | ab1670 |
| CD204 | | 1:200 | 14-905-82 |
| SDF1 | | 1 :100 | ab155090 |
| CD3 | | 1 :300 | ab11089 |
| CD8 | | 1 :200 | Homemade |
| anti-rabbit IgG | | 1:1000 | A31572 |
| anti-rabbit IgG | | 1 :500 | A32794 |
| anti-rat IgG | | 1 :500 | A48269 |
| anti-goat IgG | | 1:1000 | 705-545-003 |
| anti-mouse IgG | | 1 :500 | A32787 |
| anti-mouse IgG | | 1:1000 | A31571 |
